# An episodic burst of massive genomic rearrangements and the origin of non-marine annelids

**DOI:** 10.1101/2024.05.16.594344

**Authors:** Carlos Vargas-Chávez, Lisandra Benítez-Álvarez, Gemma I. Martínez-Redondo, Lucía Álvarez-González, Judit Salces-Ortiz, Klara Eleftheriadi, Nuria Escudero, Nadège Guiglielmoni, Jean-François Flot, Marta Novo, Aurora Ruiz-Herrera, Aoife McLysaght, Rosa Fernández

**Affiliations:** Metazoa Phylogenomics & Genome Evolution Lab, Institute of Evolutionary Biology (CSIC-UPF), Barcelona, Spain; Departament de Biologia Cel·lular, Fisiologia i Immunologia, Universitat Autònoma de Barcelona, Cerdanyola del Vallès, Spain; Genome Integrity and Instability Group, Institut de Biotecnologia i Biomedicina, Universitat Autònoma de Barcelona, Spain; Universität zu Köln, Institut für Zoologie, Cologne, Germany; Service Evolution Biologique et Ecologie, Université libre de Bruxelles (ULB), Brussels, Belgium; (IB)² – Interuniversity Institute of Bioinformatics in Brussels, Brussels, Belgium; Departamento de Biodiversidad, Ecología y Evolución, Universidad Complutense de Madrid, Spain; Department of Genetics, Trinity College Dublin, Dublin 2, Ireland

## Abstract

The genomic basis of cladogenesis and adaptive evolutionary change has intrigued biologists for decades. Here, we show that the tectonics of genome evolution in clitellates, a clade composed of most freshwater and all terrestrial species of the phylum Annelida, is characterised by extensive genome-wide scrambling that resulted in a massive loss of macrosynteny between marine annelids and clitellates. These massive rearrangements included the formation of putative neocentromeres with newly acquired transposable elements and preceded a further period of genome-wide reshaping events, potentially triggered by the loss of genes involved in genome stability and homeostasis of cell division. Notably, while these rearrangements broke short-range interactions observed between *Hox* genes in marine annelids, they were reformed as long-range interactions in clitellates. Our findings reveal extensive genomic reshaping in clitellates at both the linear (2D) and three-dimensional (3D) levels, suggesting that, unlike in other animal lineages where synteny conservation constrains structural evolution, clitellates exhibit a remarkable tolerance for chromosomal rearrangements. Our study thus suggests that the genomic landscape of Clitellata resulted from a rare burst of genomic changes that ended a long period of stability that persists across large phylogenetic distances.

## MAIN TEXT

Understanding the genomic basis of lineage origination and adaptation is key to uncovering how life diversifies and thrives in ever-changing ecosystems, providing invaluable insights into the mechanisms driving evolutionary success and the emergence of biodiversity. However, the understanding of how these processes unfold in most animal lineages has been curtailed by the lack of high-quality genomic resources, restricting comparative genomic studies to the investigation of smaller-scale events. In particular, genome-wide features and patterns, such as macrosynteny or genome architecture, have not been open to exploration.

The cornucopia of high-quality genomic resources generated by current initiatives aimed at bridging this gap is now revealing surprising information about animal evolution that can help us understand the origin of new lineages and how they adapt to changing environments, moving beyond previously known mechanisms such as point mutation or gene repertoire evolution^1^. For instance, Nakatani et al.^2^ and Simakov et al.^3^ reported chromosome-scale conservation of gene linkages (i.e. macrosynteny patterns, referred to as ancestral linkage groups, ALGs^3^) across distantly related animal phyla, revealing that many genes have remained on the same chromosome for hundreds of millions of years in animal lineages as divergent as sponges, cnidarians, molluscs and cephalochordates. This phenomenon suggests a fundamental importance of genome organisation, perhaps for gene expression regulation or other regulatory functions.

In this context, the observation of substantial genome rearrangements at the macrosyntenic level elicits great interest. Recent studies in some animal phyla, including vertebrates^2,4,5^, lepidopterans^6^, bryozoans^7^, cephalopods^8^ or tunicates^9^, demonstrated the presence of extensive rearrangements. Regarding invertebrates, in lepidopterans, Wright et al.^6^ showed that macrosynteny (inferred from lepidopteran-specific linkage groups) has remained intact across lepidopteran lineages spanning 250 My of evolution, with the exception of some lineages in which extensive reorganisation through fusion and fission was observed. Similarly, in bryozoans, Lewin et al.^7^ recently found extensive macrosyntenic rearrangements in ALGs that are otherwise highly conserved across most animal phyla, revealing that bryozoans often underwent fusion or fission. Cephalopods^8^ and tunicates^9^ also show genomic rearrangements at the level of ALGs, but to a lesser extent compared for instance to bryozoans. Notably, even in such cases of extensive chromosomal rearrangement, ALGs remain recognisable, suggesting that macrosynteny is an enduring and important feature of animal genomes.

While previous comparative studies in vertebrates have suggested some mechanisms of genome architecture remodelling, both the patterns and mechanisms of this phenomenon are still poorly understood in most invertebrate phyla. Most studies reporting genomic rearrangements focus mostly on the pattern; however, little is yet known about the mechanisms driving these rearrangements. Likewise, the architectural and functional consequences of such massive genomic changes are poorly explored in most animal phyla. Understanding these mechanisms across invertebrate phyla and their consequences for genome architecture, gene expression and function, is therefore critical for understanding the evolutionary forces that shape animal genomes across the Metazoa Tree of Life.

In this study, we generated chromosome-level genome assemblies of two earthworms from the family Hormogastridae (*Norana najaformis* and *Carpetania matritensis*), and compared them with nine annelid genomes to better understand the genomic changes leading to the colonisation of freshwater and terrestrial environments in this animal phylum. Here, we report the complete loss of macrosynteny in lineages from the same phylum (Annelida), as recently reported as well by two independent teams^10,11^, coinciding with the transition from marine to non-marine annelids (those included in the class Clitellata, comprising earthworms, leeches, potworms and their kin), to the point that ALGs are no longer recognizable. This massive genome rearrangement resulted in a complete restructuring of the genome that cannot be simply explained by typical rates of fusion and fission and rather results from genome-wide scrambling, with leeches and earthworms having among the highest rearrangement index (a metric aimed at quantifying the extent of chromosome rearrangement within genomes) among bilaterians^10^. These genomic tectonics affected chromosomal interactions, resulting in a shift towards an increase in intrachromosomal ones after chromosome mixing. We found that neocentromeres in earthworms may have evolved through the co-option of newly acquired transposable elements after scrambling, pointing to a fast centromere evolution in these clitellates. Furthermore, we identified some genetic elements that either arose or changed their position through this genome scrambling process with a putative adaptive role in the colonisation of freshwater and terrestrial environments in leeches and earthworms, respectively, indicating that the massive scrambling may have been a catalyst or facilitator to the major habitat transitions that occurred around the same time. Our findings not only shed light on the remarkable genomic transformations at the within-phylum level in annelids, resulting from a episodic burst of genomic reshaping rather than stepwise gradual genomic changes, but also suggest that clitellate genome evolution is not limited by syntenic constraints.

### Loss of macrosynteny between marine annelids and clitellates

We used long PacBio HiFi and Hi-C reads to assemble chromosome-level genomes of two earthworms of the family Hormogastridae (*N. najaformis* and *C. matritensis*) and compared them with other clitellate and marine annelid genomes available in public databases (Supplementary Data 1). We examined the macrosynteny relationships of orthologous genes across eleven chromosome-level annelid genomes (Extended Data 1). While the marine annelids displayed almost complete macrosynteny conservation with respect to the ALGs (visible as ribbons of predominantly one colour per chromosome in Fig. 1a), this relationship was shattered between marine and the clitellates (Fig. 1a,b). We explored macrosynteny conservation with the draft genome of another clitellate lineage (a potworm, *Enchytraeus crypticus*), confirming the loss of macrosynteny compared to marine annelids, and therefore indicating that this loss occurred earlier than the divergence of earthworms and leeches (Fig. 1b). After this rampant genomic reorganisation, macrosynteny in leeches and earthworms was characterised by additional massive rearrangements occurring between both groups, which are due to chromosome fusion and fission rather than genome-wide chromosome scrambling (Fig. 1c,d, Fig. 2b,c).

**Figure 1.**
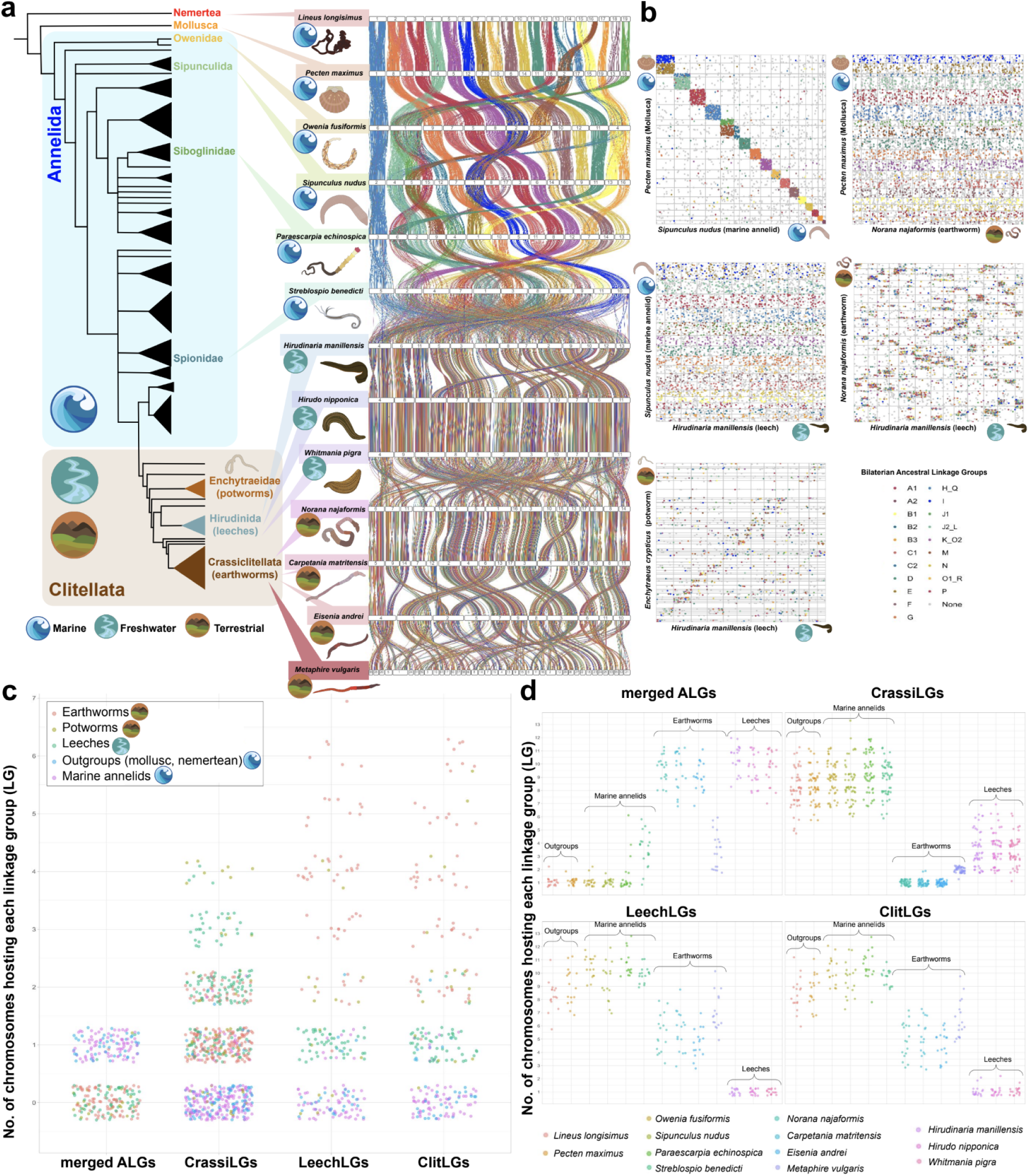
Macrosyntenic evolution of clitellates. **1a.** Left, Annelida Tree of Life showing all main lineages and the position of the species included in the present study, based on the topology reported in Capa et al. (Annelida)^95^ and Erséus et al. (Clitellata)^17^. Centre, ribbon plot of the chromosome-scale ancestral gene linkage across annelids, with a mollusc and a nemertean as outgroups. For each species, each white rectangle represents a chromosome and the number inside represents the rank order by size of that chromosome in that species. The vertical lines (ribbons) connect the orthologous genes from each genome (see Methods). Colour code follows the Bilaterian-Cnidarian-Sponge Linkage Groups (BCnS LGs) as in^3^. **1b.** Oxford dotplots of the chromosome-scale ancient gene linkage in an early-splitting marine annelid (*Sipunculus nudus*), two clitellates (the earthworm *Norana najaformis* and the enchytraeid *Enchytraeus crypticus,* draft genome) compared to a mollusc (*Pecten maximus*) and a leech (*Hirudinaria manillensis*), showing high macrosyntenic genome conservation between outgroups and marine annelids and the complete rupture of macrosynteny between marine annelids and clitellates. Vertical and horizontal lines define the border of the chromosomes of each pair of species being compared. Each dot represents an orthology relationship between genes at the x and y coordinates. Colour code is as per Fig. 1a. **1c.** Scatterplot showing the number of chromosomes from each main lineage hosting each linkage group (LG) from four datasets: merged ancestral linkage groups (mergedALGs), Crassiclitellata (CrassiLGs), Hirudinida (LeechLG) and Clitellata (ClitLGs). **1d.** Scatterplot showing the number of chromosomes from each species hosting each Linkage Group from the four datasets described in Fig. 1c. All artwork was designed for this study by Gemma I. Martínez-Redondo.

**Figure 2.**
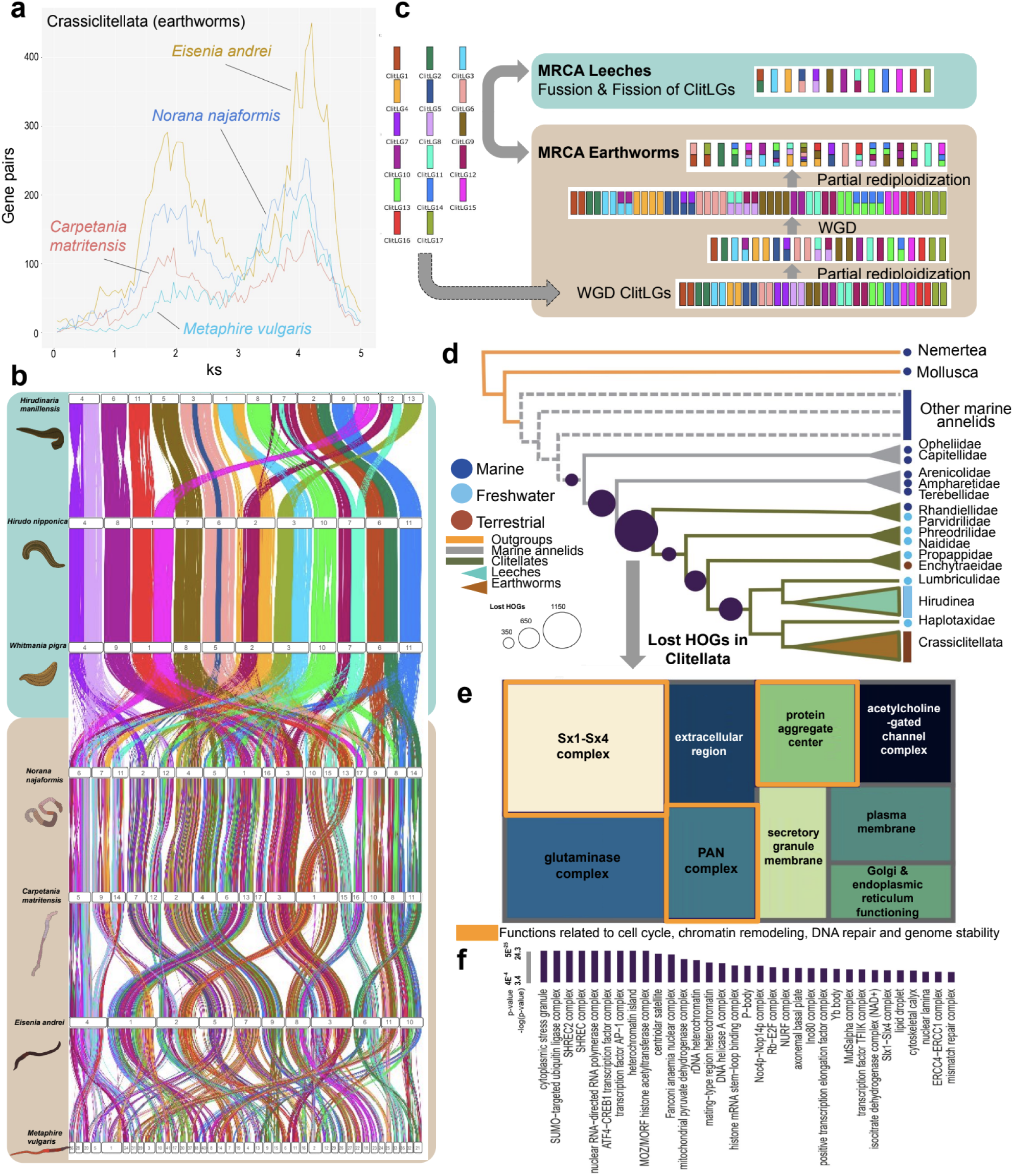
Frequent rare large-scale genomic changes among clitellates after genome-wide chromosome scrambling. **a.** Substitution-rate-adjusted mixed paralog–ortholog Ks plot for the node of Crassiclitellata (earthworms). The inferred two putative WGD events are indicated by the two prominent peaks corresponding to the Ks-based WGD age estimates. **2b.** Chromosome-scale ribbon plot showing macroevolutionary patterns of Clitellate Linkage Groups (ClitLGs) between leeches and earthworms. The vertical lines (ribbons) connect the orthologous genes from each genome (see Methods). Colour code follows the legend in Fig. 2c. **2c.** Ancestral genome reconstruction of the macroevolutionary events leading to the ancestral genome of the Most Recent Common Ancestor (MRCA) of leeches and earthworms, respectively. WGD, whole-genome duplication. **2d.** Phylostratigraphic pattern of gene loss (measured as loss of Hierarchical Orthologous Groups, HOGs) in clitellates and the surrounding nodes. Taxon families are shown at the tips (see also Supplementary Table 12). **2e.** Simplified treemap representation of putative functions enriched in lost HOGs in Clitellata, cellular component (one-sided Fisher test, p-value < 0.05). The size of the square is proportional to the p-value of the enrichment. The most general term per square is shown for simplicity (see Suppl. Mat. for further information). Orange squares comprise functions related to cell cycle, chromatin remodelling, DNA repair and genome stability. **2f.** Extended list of functions lost in Clitellata related to cell cycle, chromatin remodelling, DNA repair and genome stability (one-sided Fisher test, p-value < 0.05). Terms are shown when the p-value of the enrichment is lower than 4e^-4^ (see also Supplementary Data 4 and the manuscript’s GitHub repository).

To test the extent of chromosome mixing resulting from these extremely unusual genomic tectonics, we investigated whether the scrambling of ALGs followed a random distribution. We defined clitellate-specific linkage groups (ClitLGs hereafter) as groups of genes whose presence on the same chromosome was conserved across leeches and earthworms. We inspected these to check if they were enriched in any ALGs, and found a complete lack of statistical enrichment, thus supporting a tectonic process characterised by random genome-wide fusion-with-mixing in a process of genome atomisation (Fig. 1c,d; Supplementary Table 1). These observations thus disfavour typical models of chromosomal fusion and/or fission where parts of the constituent chromosomes are still recognizable, as occurs in most animal phyla where extended rearrangements have been reported^6,7,9^. In addition, the genomes were unalignable (i.e. alignment-based ancestral genome reconstruction recovered a highly fragmented sequence for the most recent common ancestor of clitellates and marine annelids; Supplementary Table 2), which also suggests genome-wide chromosome mixing.

To test whether the observed rearrangements could be the result of a whole-genome duplication (WGD) event followed by differential gene loss and chromosomal reshuffling, as seen in vertebrates^2,12–16^ we inferred the presence of such events through the analysis of synonymous substitutions (Ks) taking into account synteny and gene tree - species tree reconciliation methods. None of the methods supported a WGD event at Clitellata (Fig. 2a; Supplementary Data 2), therefore discarding that differential gene loss accounts for the observed genome structure. In earthworms, both methods recovered evidence of WGD. Gene tree-species tree reconciliation methods clearly supported two rounds of WGD for all earthworm species (Crassiclitellata, Fig. 2a). On the contrary, the methods based on Ks using synteny blocks as anchors provided conflicting results, pointing to different numbers of potential WGD events depending on the species(between one and two events in earthworms), and the node in which they happened varied largely (Supplementary Data 2). We further explored the stoichiometry between ClitLGs and CrassiLGs, as under a scenario of two rounds of WGD we would expect to observe a 1:4 relationship between them. While observing a higher density of 1:2 and 1:4 relationships, other relationships such as 1:3, 1:5 and 1:6 could also be observed (Fig. 1c). We argue that while there is evidence pointing to one or several rounds of WGD, the fast dynamics of macrosyntenic changes within clitellates make their robust inference challenging, and, in any case, they seem to be related to the clade of Crassiclitellata.

### Gene loss related to genome stability in clitellates

High-quality transcriptomes of all main clitellate lineages are available^17–19^, enabling the exploration of gene repertoire evolution. Given the extensive genome-wide chromosome scrambling that we have observed, we expect the generation of gene fragments that are no longer functional^20^ during the genome fragmentation process, leading to an increase in gene loss in the branches where the rearrangements occurred. Gene repertoire evolution across an extended clitellate dataset revealed that gene loss in the branch leading to Clitellata was ca. 25% higher than in the surrounding branches (Fig. 2d), consistent with the scenario of a genome atomisation event in the branch leading to clitellates as the origin of such massive rearrangements.

To test the potential functional consequences of the increased gene loss at the origin of clitellates, we investigated the putative functions of the genes lost at that branch. These were largely enriched in functions related to cell division, DNA replication and DNA repair. Examples include complexes Slx1-Slx4, GINS, MutSalpha or SHREC, all involved in DNA damage processing^21^, DNA replication initiation and progression^22^, mismatch repair^23^, or chromatin organization^24^, respectively (Fig. 2d; Supplementary Data 3, 4; Supplementary Table 3). We hypothesise that the loss of these genes could be the underlying cause of a high number of rare genome-reshaping events observed in leeches and earthworms, such as several rounds of putative WGD (Supplementary Data 2), massive genomic rearrangements between leeches and earthworms (Fig. 2b), common fission and fusion of ClitLGs in leeches (Fig. 2c) and high levels of gene loss, duplications and rearrangements in *Hox* genes in both lineages, as discussed below. Our results therefore may suggest that the presence of relaxed selective constraints (in this case, resulting in a less efficient DNA repair mechanisms) can facilitate the occurrence and fixation of genome reshuffling, as recently proposed in rodents^25^.

### Domestication of transposable elements into neocentromeres

In order to explore if clitellate genomes show a different transposable element (TE) blueprint compared to marine annelids and its potential relationship with the observed rearrangements, we investigated TE organisation and evolutionary dynamics across the 11 annelid species. TE landscapes differed considerably in clitellates and marine annelids (Fig. 3a,b; Supplementary Table 4). Enchytraeids and earthworms exhibited similar profiles, characterised by the expansion of several TE superfamilies including DNA/hAT-Charlie and a large-scale expansion of LINE/L2, the latter previously described^26^ (Fig. 3b). TE superfamily composition differed considerably in leeches compared to marine annelids and the other clitellates, with prominent expansions of DNA/hAT-Charlie, DNA/hAT-AC, DNA/hAT-Tip100 and LINE/CRE. We found one leech-exclusive TE superfamily (LINE/Dong-R4) that accounted for ca. 5% of genome coverage (Fig. 3b). Remarkably, the leech *Hirudinaria manillensis* showed a rather dissimilar pattern compared to the two other species included in this study (*Hirudo nipponia* and *Whitmania pigra)*(Fig. 3a). We investigated whether synteny breakpoints were disproportionately associated with specific repetitive sequences, but due to the massive genome scrambling, we failed to detect any significant signal indicating a higher or lower presence of specific TE superfamilies than expected from random iterations. Therefore, the role of TEs in the extreme genome scrambling observed in clitellates remains unclear.

**Figure 3.**
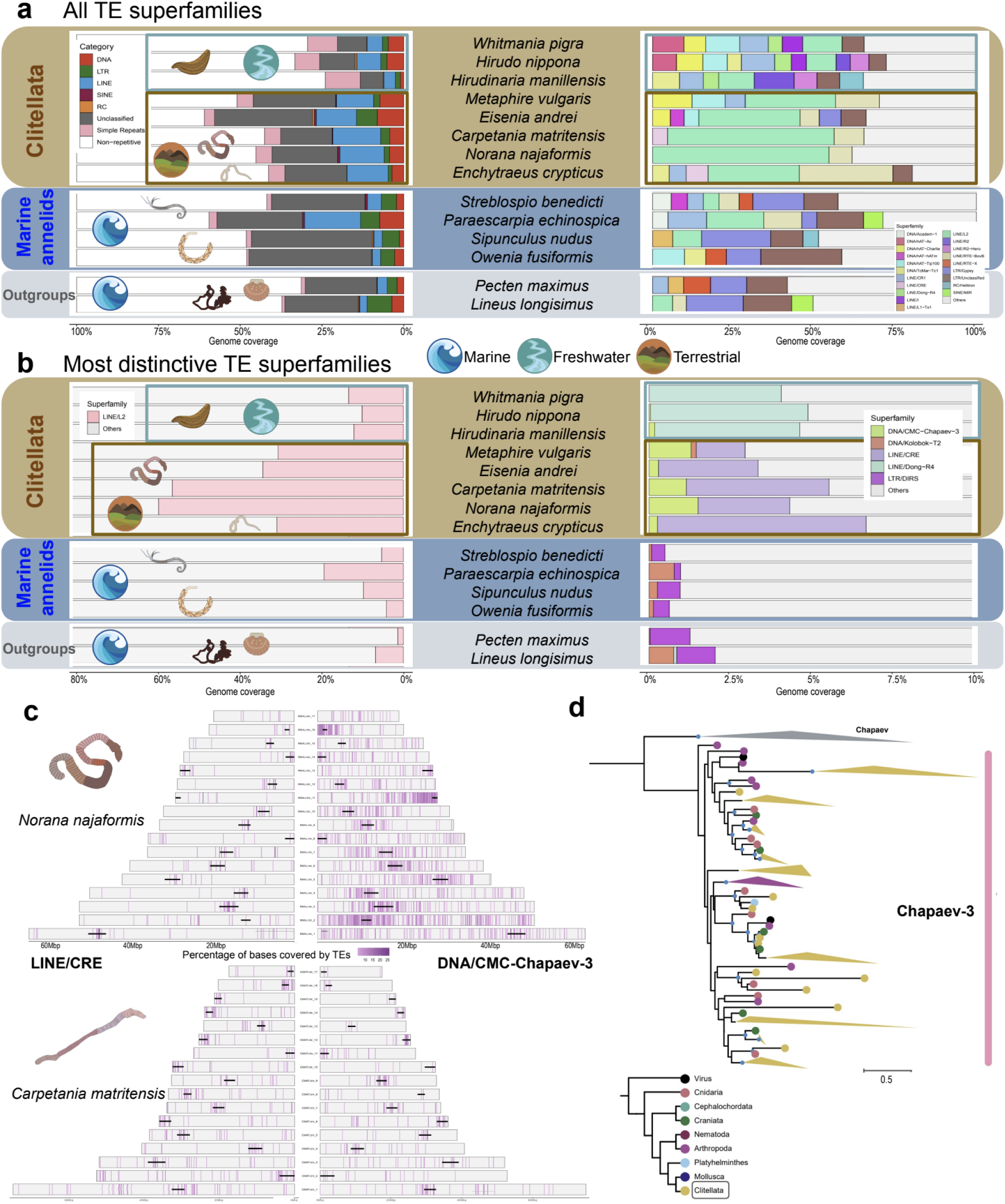
Transposable element landscape and centromere composition in clitellates. **3a.** Left, percentage of the genome of each species covered by transposable elements, divided in main categories. Right, transposable element superfamily distribution per species. The most abundant superfamilies are shown. **3b.** Left, genome coverage by the L2 superfamily. Right, genome coverage of the most qualitatively distinctive transposable element superfamilies (i.e.i.e., pattern of presence/absence in the different lineages). **3c.** Distribution of CMC-Chapaev-3 and CRE transposable element insertions in the genomes of *N. najaformis* (top) and *C. matritensis* (bottom). The genome was divided in bins of 50kb and the percentage of bases covered by members of the CMC-Chapaev-3 (right) and CRE (left) superfamilies in each bin is shown. Each horizontal bar represents a chromosome. The putative centromeres are shown with a bold black line when they could be inferred with confidence (see Methods). **4d.** Maximum likelihood phylogenetic tree of *CMC-Chapaev-3* transposase-encoding genes. Small blue dots at nodes represent clades with SH-aLRT support ≥80% and ultrafast bootstrap support ≥95%. Clitellate sequences are indicated in yellow.

Given the extent and the apparent speed of the observed genome rearrangements, we considered the effect this might have had on centromeres. To investigate whether the composition of centromeres, which are known to be monocentric in all annelids including clitellates^27^, changed after this period of massive rearrangements, we inferred the TE landscape as well as that of satellite DNA, since they both constitute centromeres^28,29^. Among the more than 100 TE superfamilies explored, only two were exclusively present in clitellates and absent in marine annelids and outgroups: the CMC-Chapaev-3 superfamily, belonging to DNA TEs, and the CRE superfamily, belonging to LINE TEs (Fig. 3b; Supplementary Table 4). Notably, these TEs were also significantly enriched in the centromeric areas in earthworms (Fisher’s exact test, p-value < 0.05; Fig. 3c). TEs from the CMC-Chapaev-3 superfamily were inferred as only present in clitellates and absent in marine annelids and leeches, and TEs from the CRE superfamily were present in enchytraeids and earthworms and absent in leeches, marine annelids and outgroups (Fig. 3b, Supplementary Table 4). Regarding the inference of satellite DNA, we failed to identify a predominant motif coinciding with the putative centromeres, and in addition, the identified ones did not coincide with the location of the centromere in clitellates, which may be indicative as well of neocentromere formation^34^.

To understand where these clitellate-exclusive TEs could have come from, we inferred a maximum-likelihood phylogenetic tree with the *CMC-Chapaev-3* TE transposase-encoding genes, including multiple species (see Methods). Earthworm transposase-encoding genes were frequently closely related to distant animal phyla such as arthropods, cnidarians or vertebrates but also viruses, indicating that they may have been acquired through horizontal gene transfer or via viral infection (Fig. 3d; Supplementary Data 5). Unlike earthworms, leech centromeres did not show enrichment of any specific TE superfamily, suggesting that centromere evolution in leeches may follow a different mechanism, potentially relying on other genomic elements for centromere function and stability.

### Reshaping of genome architecture in clitellates

A detailed comparative analysis of the genome architecture in representative annelid species (a marine annelid, *Terebella lapidaria*; an earthworm, *Norana najaformis*; and a leech, *Hirudinaria manillensis*) revealed distinct patterns of genome-wide chromosomal interactions when compared to model species (the fruit fly and chicken) (Figs. 4, 5). At the chromosomal level, annelids showed clustering of centromeres (Fig. 4a-c, center), mirroring previous observations in mosquitos^30^ and marsupials^31^. At the sub-Mbp scale, annelids exhibited compartmentalised topologically associated domains (TAD-like domains) defined by low insulator scores (Fig. 4a-c, right).

**Figure 4.**
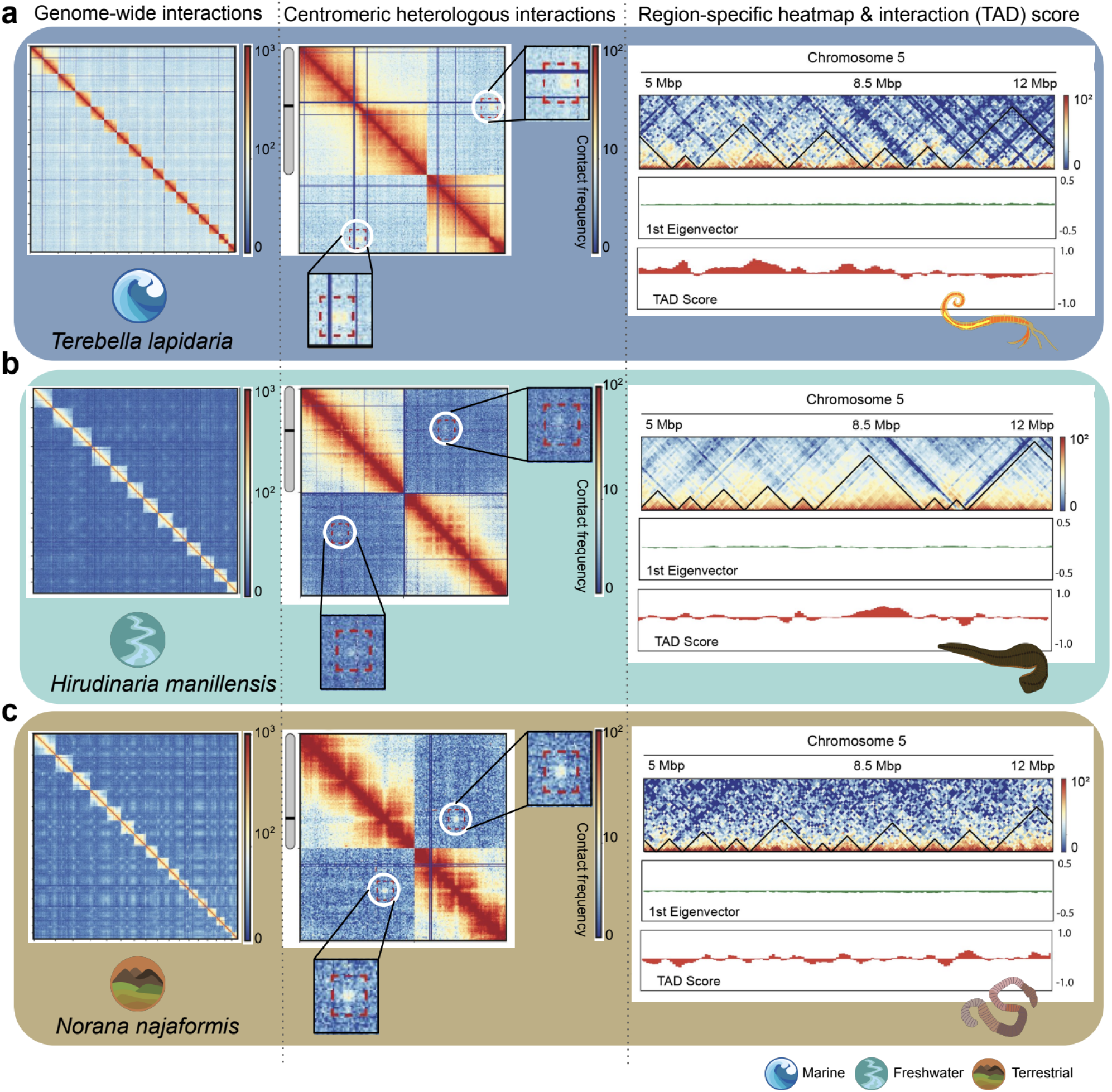
Genome architecture organisation in marine annelids and clitellates. **4a-c.** Left, whole-genome Hi-C contact maps at a 50 kbp resolution for *Terebella lapidari*a (**4a**), *Hirudinaria manillensis* (**4b**) and *Norana najaformis.* (**4c**). Center, Hi-C contact maps representing a pair of chromosomes/scaffolds depicting centromeric heterologous interactions in all three species. Right, Chromosome 5 region-specific 50 Kbp heatmaps (from 5 Mbp to 12 Mbp), depicting compartment signal (first eigenvector) and insulator (TAD) score calculated with FAN-C v0.91^84^ (see Methods), for the same three species.

To assess the robustness of the detected TAD-like domains, we recalculated them using TADbit^32^, which provides a score from 0 to 10, with 10 indicating very robust TAD-like domains (see Methods). Our results show that although at least half of the genome in annelids were compacted with strong boundary scores (>7), a significant proportion of TAD-like domains had weaker scores (<7)(Fig. 5a; Supplementary Data 6). Additionally, when creating aggregated TAD boundaries contact maps, we observed a clear decrease in interactions at TAD-like domain boundaries (Fig. 5b) Interestingly, this depletion of interactions was only enriched at the boundaries, suggesting a less organised intra-chromosomal structure. The mappability of boundary regions was consistent with, or higher than, the overall genomic mappability, indicating that the observed depletion of contacts at boundaries is not due to mapping biases but reflects genuine features of the chromatin architecture (Supplementary Data 13). Altogether, these results suggest that while annelids do exhibit TAD-like domains, their boundaries may be less well-defined, leading to a more dynamic or permissive interaction landscape within the genome.

**Figure 5.**
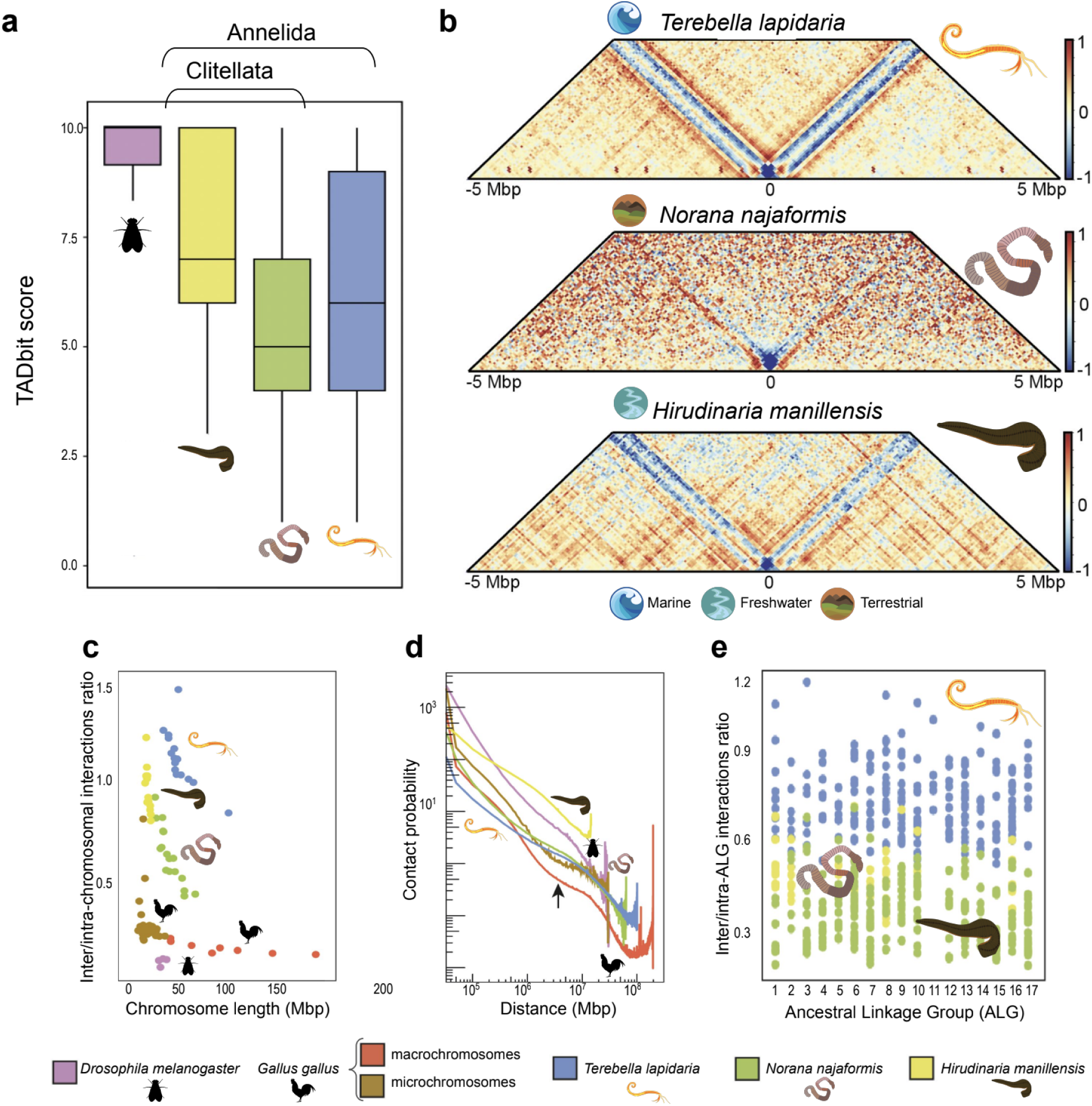
Annelid TAD-like domain organisation and inter-intra-chromosomal interactions. **5a.** Genome-wide TADbit scores per species as inferred by TADbit^32^. Box plots represent the distribution of the data, where the central line indicates the median, the box bounds correspond to the interquartile range (IQR; 25^th^ to 75^th^ percentile), and the whiskers extend to the minimum and maximum values within 1.5 times the IQR. Number of TADs analysed per species: 410 (*D. melanogaster*), 332 (*H. manillensis*), 1506 (*N. najaformis*) and 1490 (*T. lapidaria*). **5b.** Aggregated TAD plot of annelid species at 50 Kbp resolution, showing a decrease in interactions at TAD-like domain boundaries. **5c.** Inter-/intra-chromosomal interactions according to chromosome length (in Mbp) in the same three annelid species, together with chicken (*Gallus gallus*^96^) and the fruit fly (*Drosophila melanogaster*^97^). **5d.** Chromosome-specific contact probability P(s) as a function of genomic distance in *T. lapidaria*, *N. najaformis*, *Hirudinaria manillensis*, chicken and the fruit fly. **5e.** Inter-/intra-chromosomal interactions according to ancestral linkage groups (ALG) for the annelids *T. lapidaria*, *N. najaformis* and *Hirudinaria manillensis*. *[Artwork either designed explicitly for this study by Gemma I. Martínez-Redondo or retrieved from PhyloPic with a Universal Public Domain Dedication licence in the case of the <u>fly</u> and the <u>chicken</u>]*.

Remarkably, annelids showed higher inter/intra-interaction ratios per chromosome (Fig. 5c) when compared to model species, suggesting that their chromosomes are ‘floppy’ and relaxed. Additionally, annelids were heavily enriched in inter-chromosomal interactions compared to chicken and fruit fly (Fig. 5c). This difference was further supported by the analysis of distance-dependent interaction frequencies represented as curves of contact probability as a function of genomic distance, P(s) plots (Fig. 5d). The slope of the P(s) curve reflects how rapidly the contact probability decays with increasing genomic distance: a steeper slope indicates more compact chromatin folding, while a shallower slope suggests a more relaxed chromatin structure. Although the annelid curves fall between those of the fruit fly and chicken in terms of absolute values, annelids exhibit a shallower decay slope, indicating a more relaxed chromatin organization. This is consistent with the observed enrichment of inter-chromosomal interactions in annelids compared to model organisms (Fig. 5c). Such a distinct chromatin organisation suggests that annelid genomes have a more interconnected genome architecture, which may have important implications for gene regulation and genome function in these organisms.

When comparing marine annelids and clitellates, chromosome scrambling resulted in a shift in inter/intrachromosomal interactions regardless of genome size. The inter- to intra-chromosomal interactions ratio in clitellates was lower, indicating a reduction of inter-chromosomal interaction frequency and an increase of intra-chromosomal ones, a reflection of more compacted chromosomes. This pattern was maintained when comparing bilaterian ancestral linkage groups (ALGs) (Fig. 5e; Supplementary Data 7), showing that the shift towards a more compact chromatin architecture in clitellates is consistent even when considering conserved genomic regions. This suggests that despite extensive chromosomal rearrangements, the ALGs have retained a structured interaction pattern.

An exception to this pattern was the case of *Hox* genes. They showed a canonical clustered organisation in marine annelids (Fig. 6a), but in clitellates they were both duplicated and highly rearranged in different chromosomes (Fig. 6a,b; Supplementary Data 8). However, *Hox* genes from the same ancestral cluster maintained clear long-range interactions both between two separated regions of the same chromosome and between chromosomes in earthworms, indicating that they interact at the 3D level despite not being physically placed in a gene cluster (Fig. 6b-e). In leeches, however, many *Hox* genes were lost and inter-chromosomal interactions between *Hox* genes could not be detected, suggesting a deeper reshaping of the *Hox* gene repertoire (Fig. 6a).

**Figure 6.**
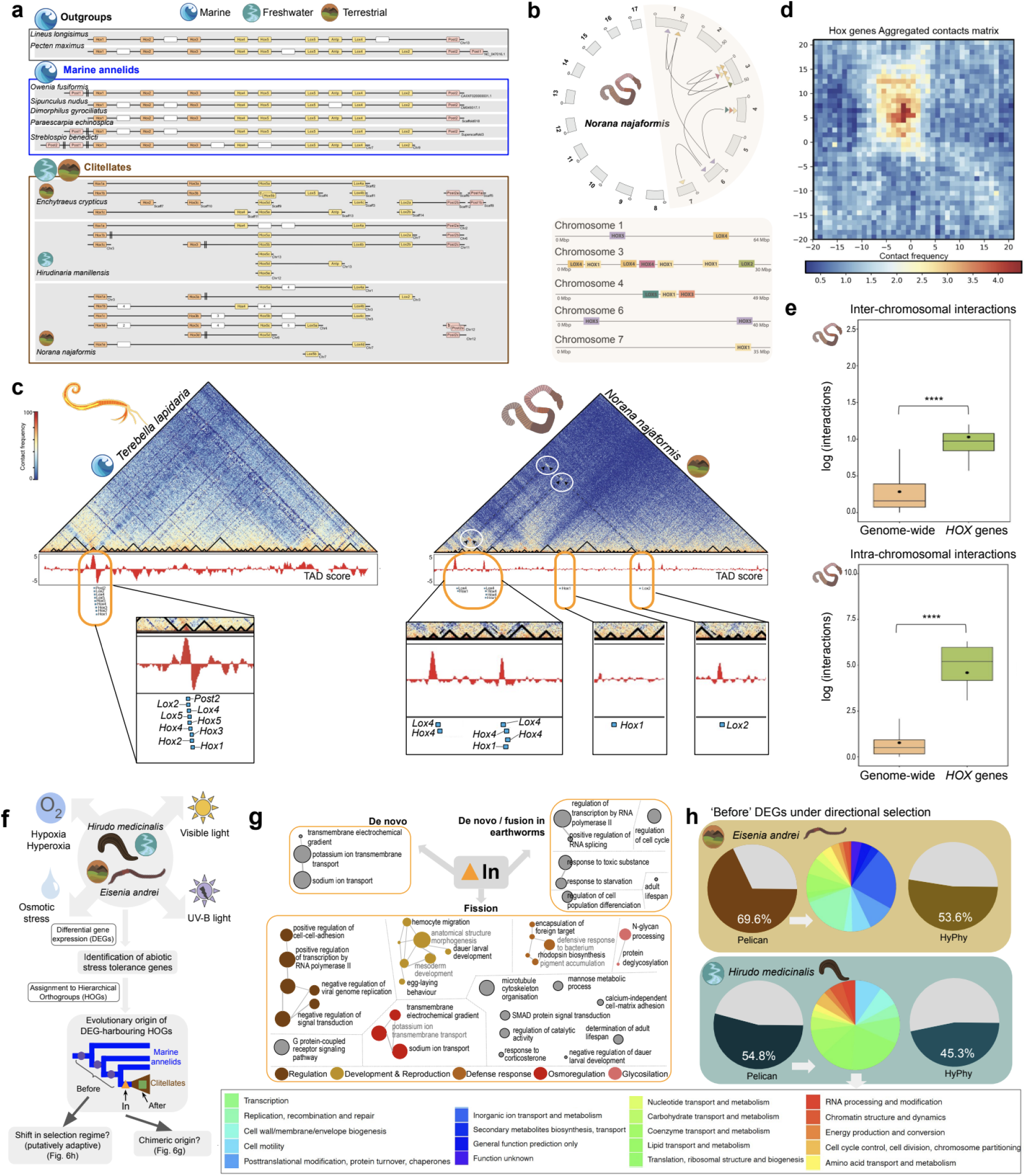
*Hox* gene repertoire and 3D interactions in annelids, and new genes involved in adaptation to freshwater and terrestrial environments in clitellates. **6a**. Representation of the main *Hox* cluster across annelids and outgroups. Rearrangements are primarily observed in clitellates. Rectangular boxes represent *Hox* genes, while chromosomes or scaffolds (in the case of the draft genome of the enchytraeid) are symbolised as distinct horizontal lines. Tandem duplications are depicted by duplicated rectangles and are named with consecutive letters. Genes not classified as a *Hox* gene are represented by empty rectangles, with the number of such genes depicted inside the triangle. Separations greater than 300kb between two non-consecutive *Hox* genes are represented by two vertical lines (||). Chromosomes are split in different lines when there are multiple copies of several genes of the *Hox* cluster in the same chromosome. **6b.** Circos plot depicting the long-range interactions involving the *Hox* genes in *N. najaformis*. **6c,d.** Chromosome-wide Hi-C map at 50 Kbp resolution of *T. lapidaria* (**6c**) and *N. najaformis* (**6d**), where clear long-range intra-chromosomal interactions between *Hox* genes can be seen in *N. najaformis* (highlighted with white circles). **6d**. *Hox* genes aggregated contact matrix is shown for *N. najaformis*. **6e.** Boxplot reflecting *Hox* genes inter-chromosomal and intra-chromosomal interactions using genome-wide interactions as reference in *N. najaformis* (two-sided t-test, ****p<0.001). Number of interactions analysed per species (shown as inter-chromosomal/intra-chromosomal): genome-wide (2783034/2227245), *Hox* genes (223/38). Boxplots show data distribution, with the central line showing the median, the box bounds the interquartile range (IQR, 25^th^–75^th^ percentile), and whiskers extend to 1.5×IQR. Outliers beyond this range are shown as points. **6f.** Schematic representation of the experimental design for the investigation of differential gene expression on *H. medicinalis* (leech) and *E. andrei* (earthworm) under abiotic stress conditions. **6g.** Putative enriched function of genes arising during chromosome scrambling (biological process). Coloured networks represent clusters of functions related ontologically; a general biological process is provided for these. **6h.** Percentage of genes under significant shifts in directional selection for both species. Results for both methods tested are shown (Pelican and HyPhy, see Methods). Main categories of DEGs significantly under directional selection in *E. andrei* and *H. medicinalis*.

### Emergence of potentially adaptive genes

Chromosome scrambling can give rise to chimeric genes resulting from the fusion of older fragments that may end up acquiring coding potential (e.g.^33,34^), allowing them to create novel proteins or regulatory functions that can help organisms adapt to new environmental challenges. To test whether chromosomal rearrangements played a role in the adaptation of clitellates to freshwater and terrestrial environments, we first explored the evolutionary origin of all hierarchical orthologous groups (HOGs) through phylostratigraphy and categorised them in three groups: (i) arising in the branches before the genome mixing (i.e. those comprised between the branch leading to clitellates and the root of the tree, containing genes from marine annelids and clitellates), which represent genes that were physically relocated during the chromosome scrambling and were therefore potentially subject to shifts in gene expression, selection, regulation or interaction with other genetic elements (hereafter ‘before’ HOGs); (ii) arising in the branch leading to Clitellata, and therefore coinciding with the genome-wide chromosome scrambling observed at that branch and comprising only clitellate sequences (hereafter referred to as ‘in’ HOGs); and (iii) arising after the genome scrambling, composed only of clitellate sequences (hereafter ‘after’ HOGs) (Extended Data 2). To test whether chromosomal rearrangements resulted in non-neutral evolution in the genes that were relocated during these events (i.e. those found in ‘before’ HOGs), we examined the selection pressures acting on these genes. Our analysis revealed that 79.6% of the ‘before’ HOGs exhibited significant shifts in directional selection (8,108 of 10,188; Supplementary Table 5). This unexpectedly high proportion sharply contrasts with the percentage of HOGs under significant shifts in directional selection in other annelid branches (which varied from 0.9% to 10.44% in three random annelid clades; see Methods and Supplementary Table 5)l. These findings suggest that the chromosomal relocations were adaptive and played a central role in the clitellate transition to freshwater and terrestrial environments.

In order to further test whether genome-wide chromosomal scrambling facilitated adaptation to freshwater and terrestrial environments, we first conducted a series of experiments aimed at identifying putative adaptive genes by analysing differential gene expression in response to abiotic conditions characteristic of freshwater and terrestrial environments in leeches (*Hirudo medicinalis*) and earthworms (*Eisenia andrei*). (Fig. 6f; Supplementary Data 9, 10 and Supplementary Tables 6, 7, 8). The number of differentially expressed coding genes (DEGs hereafter) was very similar in both species (Extended Data 2). We categorised HOGs containing DEGs in the same three groups as explained before, based on their phylostratigraphic origin (i.e. ‘before’ HOGs, ‘in’ HOGs and ‘after’ HOGs; Fig. 6f). For both species, approximately half of HOGs containing DEGs arose before the origin of Clitellata, while the other half arose after, being therefore lineage-specific (*E. andrei*: ‘before’ DEGs, 43.25%; ‘after’ DEGs, 52.59%; *H. medicinalis*: ‘before’ DEGs, 42.86%, ‘after’ DEGs, 52.04%)(Extended Data 2, Supplementary Table 9). This indicates that the gene repertoire involved in response to abiotic stress is composed of both ‘ancient’ genes (arising early in the evolutionary history of these organisms) and more recently gained, lineage-specific genes. Only 5 and 12 HOGs in *H. medicinalis* and *E. andrei* respectively, arose in the branch leading to clitellates (Extended Data 2). Remarkably, the number of DEG-harbouring HOGs in both species showed virtually no overlap (only 3 HOGs were shared; Supplementary Table 10), meaning that both species leverage a completely different gene repertoire to face environmental stress.

We next explored whether HOGs containing DEGs newly emerged in Clitellata (i.e. ‘in’ DEGs) were chimeric, as potentially expected under a scenario of chromosome shattering. Our analyses suggested that these HOGs resulted from fission of ‘ancient’ HOGs,with homology to genes encoding small heat shock proteins, transcription factors, proteins involved in arousal from lethargus in *C. elegans*, photoreception and antistatins (anticoagulants firstly discovered in leeches), among others (Fig. 6g; Supplementary Table 11). Only one HOG could be identified as originating de novo bona fide (i.e. not resulting from HOG fusion and fission), and encoded a protein putatively involved in sodium-potassium cell trafficking (Supplementary Table 11).

In order to test a potential adaptive role in relocated DEGs, we next investigated whether ‘before’ DEGs were subjected to shifts in the regime of directional selection, since such shifts could indicate that they underwent evolutionary changes to enhance fitness in response to new environmental pressures. In particular, we compared shifts in selection regimes between clitellates and non-clitellates. A high percentage of DEG-containing HOGs showed significant shifts in directional selection regimes both in earthworm and leeches (69.6% and 53.6% in *E. andrei* and 54.8% and 45.3% in *H. medicinalis* with Pelican^35^ and HyPhy^36^, respectively)(Fig. 6h). Genes with significant shifts in directional selection in the leech were involved in response to all abiotic stresses tested (visible light, UV-B light, osmotic stress, hypoxia and hyperoxia). On the contrary, more than half of the genes with significant changes in directional selection regimes in the earthworm were differentially expressed in specimens recovering after exposure to UV-B light, and therefore putatively involved in UV-induced DNA damage repair (Supplementary Table 10). The main categories based on the Clusters of Orthologous Genes (COG) database^37^ were different in both species (Fig. 6h).

### Evolutionary implications

Our study revealed massive genomic rearrangements leading to genome atomisation at the origin of the clitellates, a clade of non-marine annelids. These chromosomal tectonic shifts completely eliminated the conservation of ancestral linkage groups that are otherwise present throughout metazoans, and did so in a comparatively short evolutionary window. This extent and speed of genome restructuring is incompatible with regular models of chromosome fusion and fission, instead suggesting a process of genome-wide chromosome scrambling. While Simakov et al.^38^ and Moggioli et al.^39^ reported extensive reorganisation in the genome of a freshwater leech relative to the last common spiralian ancestor based on the analysis of draft genomes, the generation and availability of chromosome-level genomes of several clitellate lineages has revealed the timing and extent of these massive changes at the within-phylum level, potentially coinciding with the split between marine annelids and clitellates and their habitat transition towards freshwater and land. Similar findings have been reported in two recent studies^10,11^, which strengthens the robustness of the results presented here, with Lewin et al.^10^ highlighting that clitellates have among the most rearranged genomes across bilaterians.

Our results and theirs^10,11^ provide an example of a complete loss of macrosynteny genome-wide at the within-phylum level of a higher magnitude than previously seen in other animal phyla such as bryozoans^7^, cephalopodes^8^ or tunicates^9^. From a macrosynteny point of view, genome structure is much more divergent between a marine annelid and a clitellate annelid than between a marine annelid and animals as distantly related as a sponge or a mollusc, suggesting that clitellate genome evolution is not constraint by synteny. In the absence of genomic resources for other clitellate lineages to pinpoint with more precision in which branch of the Annelida Tree of Life these rearrangements may have occurred, our findings point to massive catastrophic genomic restructuring either on the branch leading to potworms, leeches and earthworms, or somewhere between the origin of clitellates and their split with marine annelids. In any case, both scenarios are consistent with a model of punctuated equilibrium^40–42^, in which a burst of genomic changes is observed in a short period after a long period of stasis (measured not in time units but as relative phylogenetic distance between lineages).

Notably, clitellates are characterised by common aneuploidy and an apparent lack of canonical cell division. Pavlíček et al.^43^ reported common aneuploidy and Robertsonian translocations in multiple species of clitellates, including earthworms, leeches and species from the families Naididade, Lumbriculidae and Branchiobdellidae, which is congruent with our findings of a peak of gene loss in clitellates enriched in functions putatively related to regulation of the cell cycle, genome stability and chromatin reshaping. Since aneuploidy is strongly associated with chromosomal and genome rearrangements and is often recognised as a direct outcome of genome instability^44–46^, our results may support the hypothesis of a single genome atomisation catastrophic event in the branch leading to clitellates resulting in the loss of genes associated to genome stability and chromatin reshaping, which may have resulted in common aneuploidy in clitellates.

The emergence of genes potentially involved in adaptation to freshwater and terrestrial environments through chromosome scrambling highlights the dynamic nature of clitellate genome architecture in response to environmental pressures. Our findings that ca. 80% of the relocated genes (’before’ HOGs and DEGs) were subject to significant shifts in directional selection provide compelling evidence that these chromosomal rearrangements were not neutral but adaptive. This suggests that the physical relocation of genes during chromosomal scrambling exposed them to new selective pressures, potentially reshaping their regulatory landscapes and optimising their functionality for the new ecological niches that clitellates encountered. Furthermore, due to their extensive nature and the high degree of selective pressure in genes relocated due to the massive chromosome scrambling, these genome-wide rearrangements may have also driven the cladogenesis of clitellates, contributing to their divergence from other annelid lineages.

Chromosome rearrangements not only facilitate the relocation of genes, which can potentially impact their regulatory landscapes and expression, but also contribute to the generation of chimeric genes, creating novel functional elements that can drive evolutionary innovation. Some of these newly arisen genes were differentially expressed, supporting their potential adaptive significance by creating novel functional elements that can drive evolutionary innovation^47,48^. The identification of both ‘ancient’ and lineage-specific DEGs suggests that adaptation to abiotic stress in Clitellata is supported by a mix of conserved and newly evolved genes. The significant shifts in directional selection observed in a substantial proportion of relocated DEGs (i.e. ‘before’ DEGs) underscore the role of chromosomal scrambling in exposing these genes to new selective pressures, likely enhancing their adaptive potential in response to terrestrial and freshwater environments. These findings align with the hypothesis that chromosomal reshuffling may act as a key evolutionary mechanism, potentially enabling organisms to refine their gene regulatory networks and functional repertoires in the face of changing environmental challenges.

One outstanding question is how this profound genome reshaping event did not result in extinction. The answer may be in the particular genome architecture of marine annelids, which seem not to be organised in compartments but rather into ‘floppy’ chromosomes with relaxed chromatin structure, which may have resulted in a high resilience to the deep genome reshaping occurring after chromosome scrambling. Further research involving a broader range of marine annelid species is necessary to confirm whether this genome architecture is indeed ancestral and contributed to the resilience observed in clitellates. Our results also suggest that the 3D genome organisation in annelids may be different to vertebrates and model organisms, since they seem to lack key structural units such as clear A/B compartments and clearly-defined TADs (Supplementary Data 12). This, together with the fact that genome evolution in clitellates may not be constrained by synteny, positions them as excellent models to further our understanding of genome evolution across the animal kingdom. A recent study found as well the lack of TAD-like domains and of syntenic constraints in platyhelminthes^49^, suggesting that our current knowledge on animal genome evolution is incomplete and that a further investigation of the genome architecture of lesser-studied invertebrates is likely to result in unexpected insights into the diversity and plasticity of genomic organisation.

All in all, our study illustrates how saltational rather than gradual changes played an important role during the evolution of an animal lineage characterised by a series of morphological and ecological innovations, providing new insights into the mode and tempo of macroevolution in animals. These results thus position clitellates as excellent model systems for further investigation into the mechanisms leading to massive genomic rearrangements and their consequences at the architectural and functional level, as well as their potential role as catalysers of cladogenesis at a macroevolutionary scale in short evolutionary windows.

## METHODS

### Specimen collection and sample processing

A detailed list of species used for each analysis is provided in Supplementary Table 12. Wet lab sample processing and sequencing is described in Extended Data 3.

### Genome assembly

A complete genome report for both species including software used for assembly, purging, scaffolding, gap filling, polishing, k-mer content assessment in the final assemblies and BUSCO completeness is available in Supplementary Data 1.

### Annotation of transposable elements

Transposable elements (TEs) were de novo identified using RepeatModeler v2.0.3^50^ with the LTRStruct flag to identify long terminal repeat (LTR) retrotransposons. The generated library of TE sequences was employed to mask the corresponding genome using RepeatMasker v4.1.2^51^. RepeatMasker was run with the following flags: “-s -a -nolow -no_is -xsmall”.

### Gene annotation

RNA-seq reads were trimmed using Cutadapt v2.9^51,52^ and assembled using Trinity v2.11.0^53^. The transcriptome assembly and the RNA-seq reads were provided as input to Funannotate v1.8.13^54^ for training using HISAT2 v2.2.1^55^ to map the reads, PASA v2.5.2^56^ and TransDecoder v5.5.0^57^. Genes were then predicted using a combination of the training outputs, Augustus v3.3.3^58^ and GeneMark-ES v4.68^59^, and processed using EVidenceModeler v1.1.1^60^.

The chromosome-level genome of *Terebella lapidaria* was explored for Hi-C analysis as described below, however gene annotation was not available in the public databases. To annotate it, we first used RepeatModeler v2.0.3^50^ to identify repeated content and RepeatMasker v4.1.2^51^ to soft mask the genome. Next, BRAKER3^61^ was used, including as input the RNA-seq data available for this species in BioProject PRJEB59381 and the OrthoDB v11^62^ input proteins corresponding to the metazoa data set, along with all of the protein sequences from annelids available in MATEdb2^63^. The annotation was isoform filtered to include only one protein per locus using AGAT^64^. Completeness of the gene annotations was evaluated using BUSCO v5.5.0^65^ with the metazoa odb_10 database.

### Inference of ancestral linkage groups

Macrosynteny between chromosome-level genomes of 11 annelid species (including the newly generated ones) plus two outgroups (a nemertean - *Lineus longissimus* - and a mollusc - *Pecten maximus*) was explored with odp v0.3.0^66^ (Extended Data 1). The Bilaterian-Cnidarian-Sponge Linkage Groups (BCnS LGs^3^) were inferred to describe the relation between these linkage groups and the chromosomes of the species in our dataset. For clarity, we merged the linkage groups that were always together in the outgroups and the marine annelids as follows: A1 comprises A1a and A1b, E comprises Ea and Eb, H_Q comprises H and Q, J2_L comprises J2 and L, K_O2 comprises K and O2, O1_R comprises O1 and R and Q comprises Qa, Qb, Qc and Qd. We referred to these linkage groups as merged linkage groups (mergedLGs). Additionally, we used odp v0.3.0^66^ to infer linkage groups specific for the leeches (named LeechLGs) using the genomes of the three leeches, and for Crassiclitellata (i.e. earthworms, named CrassiLGs) using the genomes of one earthworm per family represented (*N. najaformi*, *E. andrei* and *Metaphire vulgaris* To infer linkage groups for Clitellata (ClitLGs), we intersected the LeechLGs and the CrassiLGs. For every gene, we determined its corresponding LeechLG and CrassiLG and then evaluated if the overlap between them was significant using a Fisher test. We corrected for multiple comparisons using the Benjamini-Hochberg method and only overlaps with corrected p-values < 1e-10 were considered as candidate ClitLGs. This process was repeated for the aforementioned six species. We then compared these candidates across species to identify species-specific ClitLGs and to merge or split the candidate ClitLGs as needed. To test for the enrichment of a specific linkage group in the set of chromosomes of a species, a Fisher test was used. The list of p-values was corrected using the Benjamini-Hochberg correction method (Supplementary Table 1). The ribbon plots and the oxford dotplots were generated using custom R scripts using the ggplot2 and RIdeogram packages and the output of odp. To identify the genomic regions that corresponded to each linkage group in each species, a custom script was used (available at https://github.com/MetazoaPhylogenomicsLab/Vargas-Chavez_et_al_2024_Chromosome_shatt ering_clitellate_origins). Briefly, the region between a pair of genes corresponding to the same linkage group was considered to belong to that linkage group as long as there was not more than one gene belonging to a different linkage group between them.

### Whole genome alignment and whole-genome duplication analysis

An extended dataset of chromosome-level assemblies were aligned using progressiveCactus v2.6.0^67^ (Supplementary Table 2). Statistics for the alignment and the ancestral genomes for each node were extracted using halStats from HAL tools^68^. To identify whole-genome duplications, ksrates v1.1.3^69^ was run with default parameters and activating the collinearity analysis. *N. najaformis*. *E. andrei*, *M. vulgaris* and *H. nipponia* were used as focal species. Additionally, Tree2GD v1.0.43^70^ was run with default parameters using all annotated chromosome-level annelids plus the draft genome of *Enchytraeus crypticus*.

### Transposable element, satellite DNA and centromere identification and analysis

TEs were de novo identified within each of the available genome assemblies using RepeatModeler^50^. Putative centromeres were identified with custom scripts using their TE density. Briefly, each genome was divided into bins of 50 kb. The fraction of bases covered by TEs was calculated for each bin. These values were smoothed using the mean value of the 50 bins surrounding each bin. Next, the 10% of the bins with the highest coverage were identified and joined when adjacent. Finally, merged bins longer than 1 Mb were selected as the putative centromeres. Putative centromere positions were checked in the contact maps of the chromosome-level assemblies of some species to confirm our inference (a marine annelid - *T. lapidaria,* an earthworm - *N. najaformis*, and a leech, *Hirudinaria manillensis*), since our results support the existence of centromere interactions in annelids (see Results). Tandem repeats and satellite DNA were identified with SRF^71^. The density throughout the genome was plotted using custom R scripts available at the GitHub repository associated with this manuscript.

We assessed whether synteny breakpoints were disproportionately associated with specific repetitive sequences using the R package GenomicRanges v1.46.1^72^. We defined 10 kb flanking windows for each ALG fragment in the genome of *N. najaformis*. Due to the high level of ALG fragmentation, we almost did not retrieve any windows with clear borders (i.e. without mixing), and hence considered that clear breakpoints cannot be inferred with confidence in clitellates.

### Transposase and *Hox* genes phylogenetic inference

Phylogenetic inference of *Chapaev-3* family transposase-encoding genes and *Hox* genes is described in Extended Data 3.

### Gene repertoire evolutionary dynamics

High-quality genomic data from thirty-five annelid species and two outgroups (Nemertea and Mollusca) were used to infer gene repertoire evolutionary dynamics across the Annelida phylum (Supplementary Table 12), following the current systematic classification of the phylum^17,18,74–76^. The pipeline described in MATEdb2^63^ was used to retrieve the longest isoform for each species. Hierarchical orthologous groups (HOGs) were inferred with OMA v2.6^77^. Gene repertoire evolutionary dynamics were estimated with pyHam^78^ and curated with custom scripts. Longest isoforms in protein mode were functionally annotated with eggNOG-mapper v2^79^ and FANTASIA^80^. With the GO annotations inferred with FANTASIA, GO enrichment analysis of genes loss in each internode were calculated using a one-sided Fisher test and the elim algorithm as implemented in the topGO package^81^. To confirm that genes lost in Clitellata were enriched in pathways involved in genome stability, DNA repair, cell cycle and chromatin remodelling, the genes from those HOGs present in non-clitellate species (*Sipunculus nudus, Streblospio benedicti, Paraescarpia echinospica* and *Pecten maximus* as outgroup) were retrieved for each species and analysed in the software Reactome (https://reactome.org/, Supplementary Data 3).

### Hi-C data processing

For *N. najaformis* we used the Hi-C data newly generated in this project from a section of the midbody. The following datasets generated with the same Hi-C protocol (and thus providing comparable results) were downloaded from ENA: *T. lapidaria* (accession number ERR10851521) and *Hirudinaria manillensis* (accession number SRR15881149).

Hi-C data processing was performed using TADbit v1.0^32^. Briefly, reads were mapped in windows from 15bp to 75bp in 5bp steps using GEM3-Mapper v3^32^. Only valid pairs of mapped reads were considered to avoid noisy contacts. To achieve this, we used the following filters provided by TADbit v1.0^32^ to remove artefacts such as “self-circle,” “dangling-end,” “error,” “extra dangling-end,” “too short,” “too large,” “duplicated,” and “random breaks.” (see Supplementary Table 13). Subsequently, contact matrices were created at 50Kbp resolution and normalised to 50M contacts, following the ICE method using hicNormalize from HiCExplorer v3.7^82^. Finally, matrices were corrected and plotted using ‘hicCorrect’ and ‘hicPlotMatrix’ from HiCExplorer v3.7^82^.

### Inter-chromosome/intra-chromosome interaction ratio and averaged contact probability curves

Intra- and inter-chromosomal interactions were analysed by converting the corrected and normalised 50 Kbp matrices from h5 to interactions, using the tool ‘hicConvertMatrix’ from HiCExplorer v.3.7^82^. To obtain the inter-chromosome/intra-chromosome interaction ratio, the sum of inter-chromosomal interactions was divided by the sum of intra-chromosomal interactions per chromosome with RStudio. The correspondence between ALGs and chromosomes was used to analyse ALGs interaction distribution in the same manner.

Additionally, we generated distance-dependent interaction frequency plots, or contact probability decay curves [P(s)], to assess how contact probability decreases with increasing genomic distance. We used the corrected and normalised 50 Kbp matrices as input for the ‘hicPlotDistvsCount’s tool from HiCExplorer v3.7^82^. The P(s) curves were plotted using RStudio, with a maximum genomic distance of 1 Gbp to encompass the full range of observed intra-chromosomal interactions. Our approach follows established practices in Hi-C data analysis as described by Lieberman-Aiden et al.^83^.

### A/B compartments and TAD-like domain inference

We calculated the first eigenvector of the 50 Kbp normalised contact matrices to determine A/B compartments using the observed–expected method with the ‘fanc compartments’ tool from FAN-C v0.91^84^, with default settings. Additionally, we obtained insulator score values using the ‘fanc insulation’ tool on the normalised 50 Kbp matrices (referred to as ‘TAD score’). To validate our compartment identification, we performed a sanity check by plotting the genome-wide Pearson correlation matrix of normalised contact frequencies alongside the first eigenvector values obtained from the principal component analysis (PCA). This was accomplished using the ‘fanc plot’ tool from FAN-C v0.91^84^. We repeated the analysis with a resolution of 250 Kbp to confirm that the same pattern was observed. By overlaying the eigenvector track onto the correlation matrix heatmap, we visually assessed the correspondence between the patterns of chromatin interactions and the assigned A/B compartments, which revealed that compartments in annelids are attenuated compared to model organisms regardless of the resolution of the contact matrices (Supplementary Data 12).

TADbit v1.0^32,84^ was employed to calculate TAD boundary strength, assigning values from 1 to 10 (referred to as ‘TADbit score’), as it has proven to be more accurate in defining TADs on invertebrate species^32,85^. To ensure that the detected boundaries were not artefacts due to low mappability regions, we assessed the mappability of the genomic regions corresponding to these boundaries. We calculated mappability scores using GenMap^86^, with kmer length=30 and number of errors=2. We then compared the mappability scores of the boundary regions to the genome-wide average.

Aggregated TAD boundaries contact maps were created using the ‘hicAverageRegions’ tool from HiCExplorer v.3.7^82,86^, with TADbit TAD boundary coordinates and normalised 50 Kbp contact matrices as inputs. Plots displaying contact heatmaps, along with 1st eigenvector and insulator score tracks, were generated using the pyGenomeTracks tool (https://github.com/deeptools/pyGenomeTracks).

### Long-range interactions

Inter- and intra-chromosomal long-range contacts were identified using interaction data frames by extracting interactions that were at least twice the mean of inter- or intra-chromosomal interactions, respectively. For intra-chromosomal interactions, only contacts between loci separated by at least 2Mb were considered long-range. The long-range interactions connecting *Hox* genes were visualised using a circos plot^87^.

### Genome-wide selection analyses

To explored signatures of selection in the HOGs that arose before Clitellata and that changed their position due to the chromosome scramblingt, we selected HOGs that arose before Clitellata and that contained a minimum of 20% of the species of marine annelids and a minimum of 20% of clitellate species to ensure an adequate taxon representation (n=10,188; see also Supplementary Table 5).

To detect differential selection in protein sequence alignments at the genome-wide level in clitellates potentially facilitating the transition from marine to freshwater or terrestrial environments, we used Pelican^35^ with the option ‘pelican scan discrete’ and ‘–alphabet=AA’. In order to get gene-level predictions, we used the Gene-wise Truncated Fisher’s method (GTF) to obtain a score for all HOGs as implemented in the R package GTFisher (https://gitlab.in2p3.fr/phoogle/pelican/-/wikis/Gene-level-predictions). To calculate p-values, we considered the best 10 site-specific p-values in each alignment (k=10) and corrected for a false discovery rate (FDR) of 0.05 with the Benjamini & Hochberg (BH) method^88^. We considered as significantly under directional selection those HOGs with FDR-adjusted p-values < 0.05 (Supplementary Table 5). In order to have some expectations about the levels of selection that could be expected in other clades across the annelid evolutionary chronicle, we repeated the analysis in three annelid clades, all of them comprising species from several annelid families. Clade 1 included *Arenicola marina* (family Arenicolidae), *Neoamphitrite robusta* (family Terebellidae) and *Hypania invalida* (family Ampharetidae). Clade 2 included *Leitoscoloplos robustus* (family Orbiniidae), *Cirratulus cirratus* (family Cirratulidae) and *Paraescarpia echinospica* (family Siboglinidae). Clade 3 included *Ophryotrocha xiamen* (family Dorvilleidae), *Alitta virens* (family Nereididae), Syllis gracilis (family Syllidae), *Lepidonotopodium sp.* (family Polynoidae) and *Laetmonice cf. iocasica* (family Aphroditidae)(Fig. 2d, Supplementary Table 5 and Supplementary Table 12). For each clade, we first parsed the HOGs inferred in the category ‘before’ and containing at least 20% of marine annelids and 20% of clitellatese. From these, we selected those where all species were included in the cases of clades 1 and 2. In the case of clade 3, no HOGs included the 100% of species, and therefore we selected HOGs including at least the 80% of species. For each clade, we coded the species belonging to each marine annelid lineage as foreground and the rest of species as background. The three sets of HOGs selected for each clade were run in Pelican, with the same parameters as described above for the clitellates.

### Stress experiments and differential gene expression

Specimens from *E. andrei* and *H. medicinalis* were subjected to several types of abiotic stress related to their terrestrial/freshwater ecological niches, including exposure to visible and UV-B light, hyperoxia, hypoxia, and osmotic stress. Experiment conditions and differential gene expression analyses are detailed in Extended Data 3.

### Comparative genomics and selection analyses of genes involved in response to abiotic stress

The longest protein isoforms of the references used in differential gene expression analyses of *H. medicinalis* and *E. andrei* were included in the analysis of gene repertoire evolution as described above to infer which HOGs contained these genes. DEGs for each species and experiment were assigned to HOGs and mapped into the phylogeny to have a phylostratigraphic profile of their evolutionary origin. DEGs of *E. andrei* and *H. medicinalis* were classified in three groups based on their origin with respect to Clitellata: 1) ‘before’, 2) ‘in’, and 3) ‘after’ (i.e. they arose before the origin of Clitellata, in the branch leading to this clade - coinciding with the chromosome scrambling - or after). In the case of *H. medicinalis,* due to the lack of an available genome sequence, we identified the coordinates of orthologous sequences in HOGs inferred from the chromosome-level genome of *Hirudinaria manillensis* with corroborated high homology (reciprocal best hits in BLAST+^89^).

We tested if DEGs within HOGs that changed position during the genome-wide fusion-with-mixing (i.e, those within the ‘before’ group as described above) were subjected to directional selection. We used the software Pelican^90^. As a second selection analysis, we tested whether DEG-harbouring HOGs from the ‘before’ category were subject to positive selection. A codon based alignment for each HOG was obtained using HyPhy version 2.5.42^36^. This alignment was used to obtain a phylogenetic tree using IQTREE version 1.6.12^36,91^ selecting the MFP substitution model. Clitellate branches were tested for positive selection using aBSREL^92^. Putative function of DEGs was explored with the Cluster of Orthologous Genes (COG) database in NCBI to assign COG annotations^37^.

### Composite gene analyses

HOGs containing the differentially expressed genes that originated in Clitellata as defined in the previous section (i.e. ‘in’ group) were explored to characterise their chimeric origin based on the criteria used by Mulhair et al.^37,93^. An all vs all BLASTp of all our proteomes was performed to create a sequence similarity network with the cleanBlastp command in CompositeSearch^94^. The correspondence between the filtered HOGs and the genes was used, together with the output of cleanBlastp, as input for CompositeSearch to identify composite and component genes. Parameter values suggested in the tutorial were used (e-value of 1e-5, 30% identity, 80% coverage, maximum overlap of 20). A HOG was considered to be a composite HOG when more than half of the genes belonging to that HOG were composites. A composite HOG was considered to have originated from a fusion when most of its component HOGs were inferred to have originated before the origin of the composite HOG. Similarly, a fission origin was inferred when the age of origin of the component HOGs was younger than the age of origin of the composite HOG. In the case of component HOGs, it was considered part of a fusion when all the genes in the HOG were components, and most of the composite genes these genes were components of had an origin younger than the origin of the component HOG. Contrarily, when the origin of the composite HOG predated the origin of the component HOG, it was considered to have originated from fission.

## Data availability

The sequencing reads for the genome of *C. matritensis* are available in the ENA database under accession number PRJEB74758. Those for the genome of *N. najaformis* are available under accession number PRJEB60177. The annotated genome of *C. matritensis* is available under project PRJEB74757 and that of *N. najaformis* under PRJEB74664. The sequencing reads for the stress experiments in *H. medicinalis* and *E. andrei* are under the accession numbers PRJEB74906 and PRJEB74907 respectively. The chromosome-level genome of *Terebella lapidaria* is available in NCBI under accession number GCA_949152475.1. Data retrieved from public repositories is available under accession numbers reported in Supplementary Table 12.

## Code availability

Custom scripts are available in our GitHub repository (https://github.com/MetazoaPhylogenomicsLab/Vargas-Chavez_et_al_2024_Chromosome_shat tering_clitellate_origins; https://zenodo.org/records/15039517).

## Acknowledgements

We dedicate this manuscript to Darío J. Díaz Cosín (PhD advisor of MN and RF), who devoted his career to the study of earthworm biology and was always fascinated by them; little did we know then how truly special (from an evolutionary perspective) these creatures are. We thank Aureliano Bombarely, Rita Rebollo, Clément Goubert and Francesco Cicconardi for insightful discussions on genome rearrangements, transposable elements and tips on whole genome alignment, respectively; Pau Balart and Leandro Aristide for helping on the capture of the *Norana najaformis* specimens; Koryu Kin and Gonzalo Bercedo for kindly allowing us to use the Hypoxylab; and Natasha Tilikj and Luis Cunha for aiding in data generation for the *Carpetania matritensis* genome. GIM-R acknowledges the support of Secretaria d’Universitats i Recerca del Departament d’Empresa i Coneixement de la Generalitat de Catalunya and ESF Investing in your future (grant 2021 FI_B 00476). LA-G was supported by an FPI predoctoral fellowship from the Ministry of Economy and Competitiveness (PRE-2018-083257). NG was supported by the European Union’s Horizon 2020 research and innovation programme under Marie Skłodowska-Curie grant agreement 764840 to JFF (ITN IGNITE; www.itn-ignite.eu) as well as by a Deutsche Forschungsgemeinschaft (DFG) grant (458953049). MN acknowledges support from Ramón y Cajal fellowship (RYC2018-024654-I) and by Grant PGC2018-094112-A-I00 (which provided funding for the genome of *C. matritensis*) both from MCIN/AEI/10.13039/501100011033 and by “ESF: Investing in your future” and “ERDF: A way of making Europe” respectively. ARH acknowledges support from the Spanish Ministry of Science and Innovation (PID2020-112557GB-I00) funded by AEI/10.13039/501100011033, the Secretaria d’Universitats i Recerca del Departament d’Economia i Coneixement de la Generalitat de Catalunya (AGAUR 2021-SGR00122) and the Catalan Institution for Research and Advanced Studies (ICREA). AMcL was supported by funding from the European Research Council grant agreement 771419. RF acknowledges support from the following sources of funding: Ramón y Cajal fellowship (grant agreement no. RYC2017-22492 funded by MCIN/AEI /10.13039/501100011033 and ESF ‘Investing in your future’), the European Research Council (this project has received funding from the European Research Council (ERC) under the European’s Union’s Horizon 2020 research and innovation programme (grant agreement no. 948281)), the Catalan Biogenome Project (which provided funding for sequencing the genome of *N. najaformis*), the Secretaria d’Universitats i Recerca del Departament d’Economia i Coneixement de la Generalitat de Catalunya (AGAUR 2021-SGR00420) and the OSCARS project, which has received funding from the European Commission’s Horizon Europe Research and Innovation programme under grant agreement no. 101129751. We also thank Centro de Supercomputación de Galicia and CSIC for access to computer resources (CESGA and DRAGO respectively).

## Author contributions

CV-C and RF designed the study and coordinated data analysis. CV-C conducted analysis on macrosynteny, transposable element evolution and satellite repeat identification, whole-genome duplication and ancestral genome reconstructions. LB-A carried out analyses on gene repertoire evolution. GIM-R performed composite gene analysis and transposase and *Hox* gene phylogenies. KE conducted stress experiments and differential gene expression analysis. LA-G and AR-H led analysis of genome architecture. JS-O and NE conducted wet lab experiments and stress experiments. NG and CV-C led genome assembly and annotation. JFF assisted with genome assembly. MN generated new data for genome assembly. CV-C, LB-A, GIM-R, KE, JS-O, LA-G, AR-H, AMcL and RF interpreted the data and discussed results. RF wrote the initial version of the manuscript, with input from all authors. RF provided resources and supervised the study. All authors revised and approved the final version of the manuscript.

